# Pro-tumor and prothrombotic activities of hepsin in colorectal cancer cells and suppression by venetoclax

**DOI:** 10.1101/2022.10.01.510038

**Authors:** Maria Carmen Rodenas, Julia Peñas-Martínez, Irene Pardo-Sánchez, David Zaragoza-Huesca, Carmen Ortega-Sabater, Jorge Peña-García, Salvador Espín, Guillermo Ricote, Sofía Montenegro, Francisco Ayala de la Peña, Ginés Luengo-Gil, Andrés Nieto, Francisco García-Molina, Vicente Vicente, Francesco Bernardi, Maria Luisa Lozano, Victoriano Mulero, Horacio Pérez-Sánchez, Alberto Carmona-Bayonas, Irene Martínez-Martínez

## Abstract

Hepsin is a type II transmembrane serine protease whose expression has been linked to greater tumorigenicity and worse prognosis in different tumors such as prostate and gastric cancer. Recently, our group described hepsin expression in the primary biopsy as a potential biomarker of thrombosis and metastasis in localized colorectal cancer patients. Here we explored the role of hepsin in this tumor. Hepsin overexpression increased Caco-2 cell migration and invasion, higher phosphorylation of Erk1/2 and STAT3 and led to more thrombin generation in plasma. Indeed, our study revealed higher plasma levels of hepsin in metastatic colorectal cancer patients, which was associated with a greater tendency toward thrombosis. By virtual screening of a FDA-approved drug library, we identified venetoclax as a potent hepsin inhibitor, reducing the metastatic and prothrombotic phenotype of Caco-2 cells, but not of other colorectal cancer cells without hepsin expression. Interestingly, pre-treating Caco-2 cells overexpressing hepsin with venetoclax reduced its *in vivo* invasiveness. Taken together, our results demonstrate that elevated hepsin levels correlate with a more aggressive and prothrombotic tumor phenotype. Likewise, they evidence an antitumor role for venetoclax as a hepsin inhibitor. This lays the groundwork for molecular targeted therapy for colorectal cancer.

## 1. Introduction

Proteolysis in tumors’ microenvironment confers an adaptive advantage to emerging tumors, through their capacity to remodel the extracellular matrix and orchestrate various processes of angiogenesis, invasion, and metastasis. Multiple proteases dysregulate in tumor cells to foster progression and spread [1]. However, not all tumors develop the same strategies nor do they activate the expression of the same proteases to sustain tumorigenic processes. Notably, type II transmembrane serine proteases (TTSPs) comprise a family implicated in these processes [2]. Hepsin is one such TTSP and its expression has been linked to greater tumorigenicity in different types of cancer, such as breast, ovary, and prostate [3–5], and with worse prognosis in gastric cancer [6]. Hepsin is a key protease for Ras-dependent tumorigenesis, affecting epithelial cohesion and basement membrane integrity [7]. This seems to be relevant in tumors with a high prevalence of Ras mutations, such as colorectal cancer (CRC) [8]. In fact, hepsin is a serum marker capable of distinguishing between localized and metastatic CRC [9].

Nevertheless, hepsin’s actions on tumorigenesis are multifaceted and, paradoxically, at times even serves an antitumor role [10–13]. Thus, in different contexts, high hepsin levels have been associated with well-differentiated tumors, antitumor activities, or better prognosis. This is known as ‘the hepsin paradox’ [11,14,15]. Our group has found similar results in the biopsies of CRC patients. Thus, in metastatic individuals, low hepsin expression conducted to poorly differentiated tumors and a larger number of distant organs affected [13]. While the biological substrate has been elucidated only in part, one study conducted in prostate cancer xenografts suggests that fine-tuning can be critical, such that excessive hepsin proteolytic activity would decrease tumor cell viability, whereas moderate expression would conserve protumorigenic activity without excessive proteolytic toxicity [16]. Furthermore, hepsin has another peculiarity – it activates FVII, the factor that initiates the coagulation cascade. Hepsin’s involvement in mechanisms of thrombogenesis have been proven *in vitro* and in zebrafish models [17,18]. Recently, we have shown that hepsin from primary tumor was also a potential biomarker of thrombotic risk in localized CRC patients [13]. These observations have prompted our group to contemplate hepsin’s contribution, as a member of the TTSP family, to the hypercoagulable state of malignancy, together with other clinical and biological risk factors that come together in the cancer patients and that ultimately establish the hemostatic system activation [19,20].

In this work, we have sought to examine the functional effect of hepsin expression on colon tumor cell proliferation, migration, and invasion processes. Additionally, we have investigated the influence of hepsin levels on plasma hypercoagulability, with a view to its possible functional mechanism. Since hepsin is a potential therapeutic target in CRC, we have performed a Virtual Screening (VS) for the search of inhibitors, to subsequently characterize the antimigratory, anti-invasive, and anti-thrombotic potential of the selected compounds by means of *in vitro* and *in vivo* assays.

## 2. Materials and Methods

### 2.1. Ethics Statement

The study was conducted in accordance with the Good Clinical Practice guidelines and the Declaration of Helsinki. It was approved by the Ethics Committee of the Morales Meseguer University Hospital (EST: 07/15). All participants who were still alive at the time of data collection provided written and signed informed consent.

The experiments complied with the Guidelines of the Council of the European Union (Directive 2010/63 / EU) and RD 53/2013 of Spain. The experiments and procedures were carried out as approved by the Counseling of Water, Agriculture, Livestock and Fisheries of the Autonomous Community of the Region of Murcia (CARM authorization number # A13180602).

### 2.2. Samples and patient characteristics

Seventy-three patients were recruited with localized or advanced CRC, treated between 2012 and 2021 at Hospital Universitario Morales Meseguer. Subjects were recruited directly by the oncologist at their first appointment or during chemotherapy. The baseline characteristics of these patients are summarized in **Table 1**. All participants with advanced tumors had active disease at the outset. KRAS mutations were evaluated by real-time PCR with the Idylla system (Biocartis, Belgium).

**Table 1.** Baseline patient characteristics.

| **Characteristics** | **N (%)** |
| --- | --- |
| Age, median (range) | 63 (39-85) |
| Sex, male | 49 (67.1) |
| TNM stage |  |
| Localized | 20 (27.3) |
| Advanced | 53 (72.6) |
| Line of therapy |  |
| 1<sup>st</sup> line | 22 (30.1) |
| 2<sup>nd</sup> line | 14 (19.1) |
| 3<sup>th</sup> line | 10 (13.6) |
| 4<sup>th</sup> line | 5 (6.8) |
| Time from diagnosis of metastasis to hepsin sampling, median (range) | 4.2 (0-34) |
| Location of the primary tumor * |  |
| Right colon | 28 (38.5) |
| Transverse colon | 2 (2.6) |
| Left colon | 31 (42.4) |
| Rectum | 12 (16.4) |
| Location of metastases |  |
| Liver | 41 (56.1) |
| Lung | 20 (27.3) |
| Lymph nodes | 18 (24.6) |
| Bone | 4 (5.4) |
| Peritoneal | 19 (26.0) |
| Surgery of the primary tumor | 58 (79.4) |
| Histological grade |  |
| Grade 1 | 20 (28.7) |
| Grade 2 | 38 (52.0) |
| Grade 3 | 9 (12.3) |
| Unknown | 5 (6.8) |
| KRAS |  |
| Mutant | 22 (30.1) |
| Wild-type | 31 (42.4) |
| Not performed | 20 (27.3) |
| Chemotherapy regimen when hepsin was obtained * |  |
| FOLFOX-based | 11 (15.0) |
| Irinotecan-based | 9 (12.3) |
| Anti-angiogenic | 13 (17.8) |
| Anti-EGFR | 1 (1.3) |
| Regorafenib | 2 (2.7) |
| None | 44 (60.2) |

Abbreviations: TNM= tumor, node, metastases; KRAS= Kirsten Rat Sarcoma Viral Oncogene Homologue; FOLFOX= oxaliplatin-5-fluorouracil, EGFR= epidermal growth factor receptor.
\* Not mutually exclusive

### 2.3. Plasma hepsin determination

An *in vitro*, enzyme-linked, immunosorbent assay (RayBiotech, Norcross, Georgia) was performed to quantify human plasma hepsin in both patients with localized and those with metastatic CRC according to manufacturer’s instructions. Briefly, 1:2 dilution of plasma samples (n□=□20 localized and n=53 metastatic samples) were added to flat-bottomed 96-well plate precoated with a specific anti-human hepsin antibody, followed by a biotinylated anti-human hepsin antibody. Next, HRP-conjugated streptavidin (a biotinylated antibody) was added to the wells. Finally, TMB chromogen was added and absorbance was read at 450 nm using a plate reader (Biotek, Agilent).

### 2.4. Cell culture

Cells were cultivated using standard conditions (37°C, 5% CO2, >95% humidity). Caco-2 cell line (ATCC® HTB-37™), DLD1 and HCT-116 were purchased from American Type Culture Collection and authenticated as previously described [21]. Cells were cultivated in MEM, RPMI *(Roswell Park Memorial Institute)* and McCoy’s 5a media (Gibco-Thermo Scientific, Barcelona, Spain), respectively, containing 1% GlutaMAX (Gibco-Thermo Scientific, Barcelona, Spain), 20% fetal bovine serum (ATCC® 30-2020™, Gibco-Thermo Scientific, Barcelona, Spain) and penicillin/streptomycin (10000 U/ml, Gibco-Thermo Scientific, Barcelona, Spain), respectively. Cells were routinely checked for mycoplasma contamination.

### 2.5. Cell transfection

The plasmid DNA pCMV6-AC and the transfection control pCMV-MIR were purchased from OriGene (Maryland, EEUU). Caco-2 cells were plated (10·10^6^ cells) in 500μl of the Ingenio electroporation solution (Mirus Vio, Wisconsin, USA). The transfected cells were selected using the antibiotic geneticin/G418 (Gibco-Thermo Scientific, Spain) for 10 days. Post selection, a single cell was plated in each well of a 96-well plate to obtain a single clone.

### 2.6 Hepsin silencing

Caco-2 cells were grown in a humidified atmosphere with a 5% CO_2_ concentration. ON-TARGETplus SMARTpool siRNAs against *HPN* and control siRNA were obtained from Dharmacon (GE Dharmacon, Barcelona, Spain). siRNA transfections were performed using PepMute transfection reagent (SignaGen, MD, USA). Silencing rates were approximately 80% after two days of transfection (**Figure S1**).

### 2.7. RT-qPCR

Total RNA was obtained from Caco-2 (cells transfected with pCMV-MIR), Caco-2-HPN (HPN-overexpressing cells transfected with pCMV6-AC) and silenced cells, and from DLD1 and HCT-116 using Trizol® Reagent (Invitrogen, Carlsbad, CA). NanoDrop spectrophotometer (Thermo Scientific, Wilmington, DE) was used to determine RNA concentration and 260/280 ratio. From the total RNA, a reverse transcription of a 100-ng sample to cDNA was performed (SuperScript First Strand, Invitrogen, Madrid, Spain). PCR reactions in triplicates were performed using TaqMan® Gene Expression probes Hepsin (HPN): hs01056332_m1; β-Actin (ACTB): hs01060665_g1 on a LC480 Real Time PCR (Roche, Madrid, Spain). ACTB expression was used as the endogenous reference control using the comparative cycle threshold method, Ct method (2^−ΔΔCt^).

### 2.8. Western blot analysis

Cell lysates were subjected to 10% SDS PAGE and subsequently transferred to nitrocellulose membranes. Detection of proteins was performed by using primary rabbit anti-human hepsin antibody (Sigma Aldrich, Steinheim, Germany), pAKT (Invitrogen, Madrid, Spain), pERK1/2 (9101S, Cell Signaling Technologies, Werfen, Barcelona, Spain), pSTAT3^(Tyr705)^ (9145, Cell signaling Technologies, Werfen, Barcelona, Spain). Secondary IgG antibodies were horseradish peroxidase-coupled and visualized by the ECL kit detection. Protein expression of β-actin (Sigma Aldrich, Steinheim, Germany) was used as the endogenous reference control.

### 2.9. Wound healing assays

Caco-2 and Caco-2-HPN were grown as confluents and wounded by removing a 300–500 μm-wide cell strip through the well with a standard 200 μl pipet tip. Wounded monolayers were washed twice to take out non-adherent cells. The following three conditions were studied in Caco-2 cells: 1) basal hepsin, 2) overexpression, and 3) HPN-silenced. Wound healing was quantified after 72 h using Fiji-ImageJ software as previously described [21].

### 2.10. Degradation of gelatin coated coverslips

Gelatin matrix was prepared by mixing 0.2% gelatin and rhodamine (Invitrogen, Life Technologies, Madrid, Spain), as previously described [22]. Then, the coverslips were coated with the gelatin mixture, fixed with 0.5% glutaraldehyde for 15 min, and then washed with PBS. Caco-2 cells were cultured on these coated coverslips for 72 h. Immunofluorescence analysis was performed after fixing the cells with 3.7% formaldehyde and incubation with phalloidin (Sigma-Aldrich) and ProLong Gold Antifade medium with DAPI (ThermoFisher). Images were taken with a confocal spectral scanning microscope SP8 LEICA, analyzed with Fiji-ImageJ and GIMP software, and processed with Adobe Photoshop. Degrading gelatin cells were quantified as previously described [22].

### 2.11. Flow cytometry cell cycle and proliferation assays

Cells were cultured and when reached an exponential rate (5×10^5^ cells/ml) were seeded in 6-well plates for further analysis. The cell cycle was examined after treating the cells with RNAse and with propidium iodide (Invitrogen). To evaluate the proliferative activity of Caco-2 cells, 5-ethynyl-2’-deoxyuridine (EdU) (Thermo Fisher Scientific, Barcelona, Spain) was added for 48h. Cancer cells aliquots were then used to establish the percentage of positive cells; i.e., EdU+, detected by fluorescent-azide coupling reaction with EdU (Click-iT; Thermo Fisher Scientific, Barcelona, Spain).

### 2.12. Thrombin generation assay

Venous blood was obtained from 20 healthy donors not on any medication. Blood was drawn from the antecubital vein into non-siliconized Vacutainer® tubes containing 3.8% buffered sodium citrate (Becton-Dickinson, Rutherford, NJ). The tubes were centrifuged for 15 min at 2000 *g* at room temperature; the isolated plasma was then stored at −70°C until analysis. Platelet poor plasma was prepared as previously described [23] and kept at −80°C.

Caco-2 and Caco-2-HPN cells were cultured as confluent monolayers in 6-well plates and incubated with 1 ml of plasma for 3 h at 37°C and 5% CO2. Plasma was recovered and centrifuged for 5 min at 200 g at room temperature. Thrombin generation assay-calibrated automated thrombogram (TAG-CAT) was conducted as previously described [24] (Diagnostica Stago). Samples were processed in duplicate. The following thrombogram parameters were evaluated: (a) lag-time; (b) time to peak (ttPeak); (c) peak; (d) mean rate index (MRI); and (e) endogenous thrombin potential (ETP) [24].

### 2.13. Virtual Screening

In order to find novel and safe hepsin inhibitors, we performed a VS based on the molecular docking technique [25] with Autodock Vina, [26] processing the hepsin crystallographic structure with PDB code 1P57 against the DrugBank (https://go.drugbank.com) database of compounds (version 5.0; of 9591 compounds, including 2037 approved by the American FDA, 96 nutraceuticals, and 6000 experimental) and focusing the docking process on its catalytic site with residues HIS57, ASP102, and SER195.

### 2.14. Effect of selected drugs on hepsin activity, cell migration, invasion, proliferation, cell cycle and thrombin generation assays

Hepsin (0.05 μM) (R&D Systems, Madrid, Spain) was incubated in 50 mM Tris-HCl buffer, pH 9 buffer, with 200 μM BOC-Gln-Arg-Arg-AMC and the emitted fluorescence was recorded for 5 minutes. To test the effect of selected drugs, the same reaction was carried out, but following incubation with hepsin for 1 h at 37°C. IC50 was calculated after registering hepsin activity at different drug concentrations.

Cells were incubated in the presence and absence of 1.88 μM venetoclax to evaluate migration, invasion proliferation, cell cycle, and thrombin generation in the same conditions as described above.

### 2.15. Larval xenotransplantion assays

Wild-type zebrafish (*Danio rerio* H. Cypriniformes, Cyprinidae) were obtained from the Zebrafish International Resource Center (ZIRC, Oregon, USA). The transparent *roy^a9/a9^*; *nacre^w2/w2^* (casper) was previously described [27]. Fish were mated, staged, raised and processed as described in the zebrafish handbook [28]. Fertilized zebrafish eggs were obtained from natural spawning of fish and kept in our facilities following standard husbandry practices. The animals were kept in a 12 h light/dark cycle at 28°C. Zebrafish larvae were anesthetized as previously described [29].

Caco-2-HPN cells were cultured in presence and absence of 1.88 μM venetoclax for 48 h, then disaggregated and labelled with 1,1’-di-octa-decyl-3,3,3’,3’-tetra-methyl-indo-carbo-cya-nine perchlorate (DiI, ThermoFisher) and finally resuspended in a buffer containing 5% FBS in PBS. Injection of cells in embryos was conducted as previously described [30,31] and larvae were scored for cells dissemination by fluorescence microscopy. Caco-2 and Caco-2-HPN cell invasion score was calculated as previously reported [30,31].

### 2.16. Statistical analysis

The Kaplan-Meier method was used to estimate survival, while the Aalen-Johansen estimator evaluated the cumulative incidence. The analyses were performed by means of the log-transformation of the hepsin values. The association between hepsin (continuous, log-transformed values) and thrombosis was examined using Fine-Gray regression. Furthermore, the third quartile (Q3) of the hepsin concentration (log-transformed) was used to produce a stratified description of its association with survival endpoints. Hepsin distributions were compared by the Wilcoxon test. Survival was defined from the date of hepsin analysis until demise or last follows up. Time-to-thrombosis was defined as the time elapsed between the date of hepsin extraction until thrombosis, censoring event-free subjects and factoring in death as a competing event. The analyses were executed with R version 4.05 [32].

## 3. Results

### 3.1. Hepsin levels in plasma of CRC patients

We first evaluated hepsin by ELISA in samples from 73 subjects with CRC. Patient baseline characteristics are shown in **Table 1**. Of them, 73% had advanced tumors (n=53), whereas 27% (n=20) had localized neoplasms. The mean natural logarithm of hepsin concentration was greater in advanced *vs* localized tumors (7.4 vs 6.9, p<0.001) (pg/mL) (**Figure 1A**). In advanced tumors (n=53), the mean logarithm of hepsin concentration was 7.2 vs 7.7 pg/mL in KRAS native and mutated tumors, respectively (p=0.056) (**Figure 1B**). At the time of analysis of patients with advanced cancer, 5 thrombotic events had been detected, which comprises a cumulative incidence (CI) of 7.0% (95% CI, 1.6-17.7).

**Figure 1.**
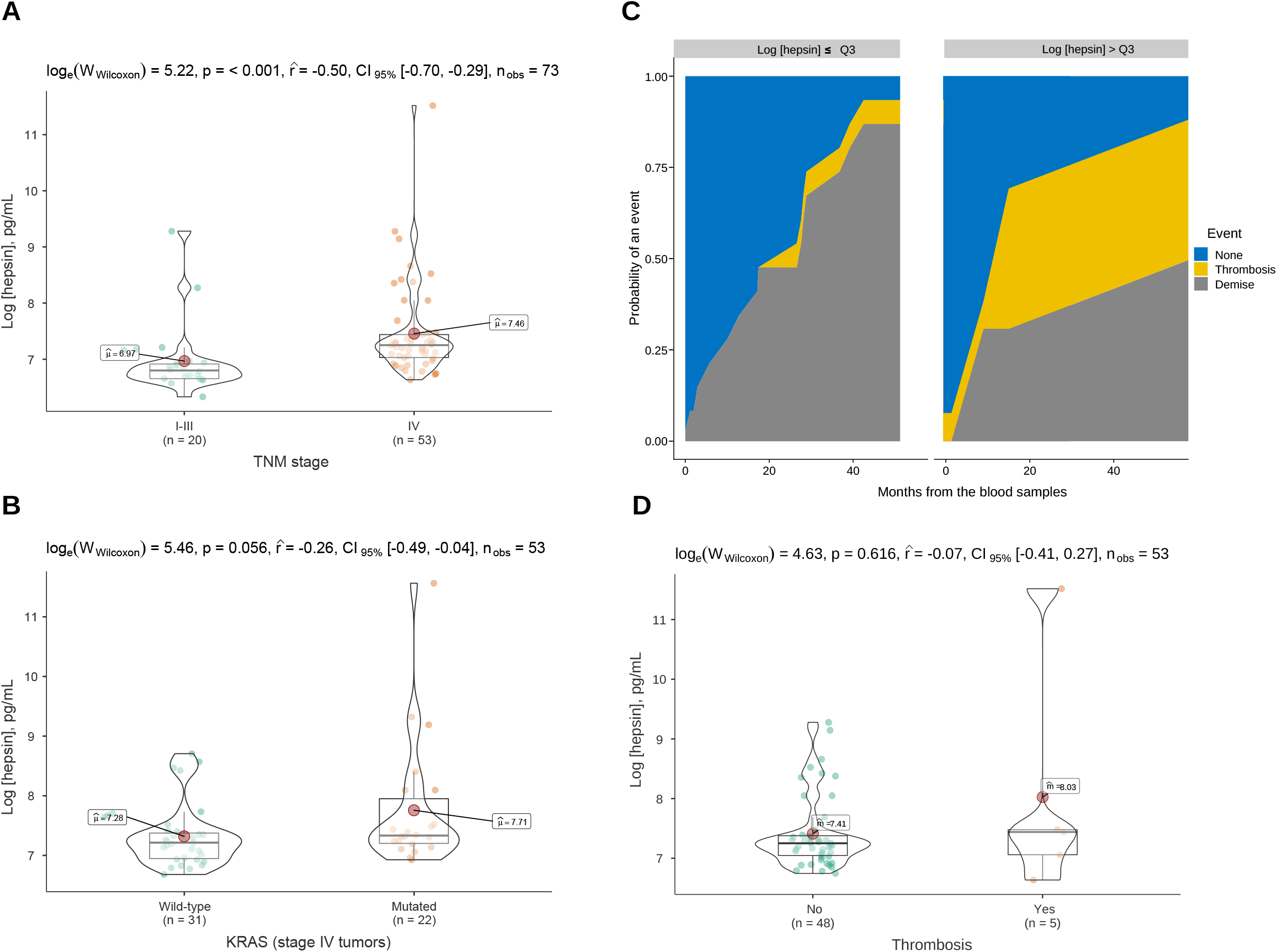
Hepsin levels in plasma of colorectal cancer patients and association with tumor stage and thrombosis. Violin boxplot showing the association between the logarithm of the hepsin concentration and TNM stage (A); and the presence of KRAS mutations in advanced cancer patients (B). C) Cumulative incidence functions for death and thrombosis among advanced cancer patients. The third quartile (Q3) of the hepsin level distribution was used to stratify the curves. D) Violin box-plots showing the log hepsin concentration in advanced cancer patients with or without thrombosis.

The subjects with a logarithm of hepsin concentration > third quartile (Q3, 7.4 pg /mL) had a 24-month CI of 34.7% (95% CI, 0.2-86) compared to 0 events in subjects with levels ≤Q3 (p-value 0.036, Gray test) (**Figure 1C**), although the result is subject to uncertainty due to the low number of events and the effect of an influential observation (**Figure 1D**). In the Fine-Gray regression, continuous hepsin (log-transformed) was associated with more thrombosis with a subhazard ratio of 2.12 (95% CI, 1.50-2.98). At present, it cannot be concluded that hepsin levels are linked to overall survival (**Figure S2**).

### 3.2. Hepsin overexpression in Caco-2 cells

The remaining experiments were conducted with Caco-2 cells. Basal hepsin expression is low in Caco-2. To boost expression, we generated clones that overexpress hepsin by means of stable transfection with the hepsin gene (HPN). As illustrated in **Figure 2A**, Caco-2-HPN exhibited a 4.4-fold increase in levels mRNA compared to the cells with basal expression, which was correlated with the protein’s effective increase in expression (**Figure 2B**).

**Figure 2.**
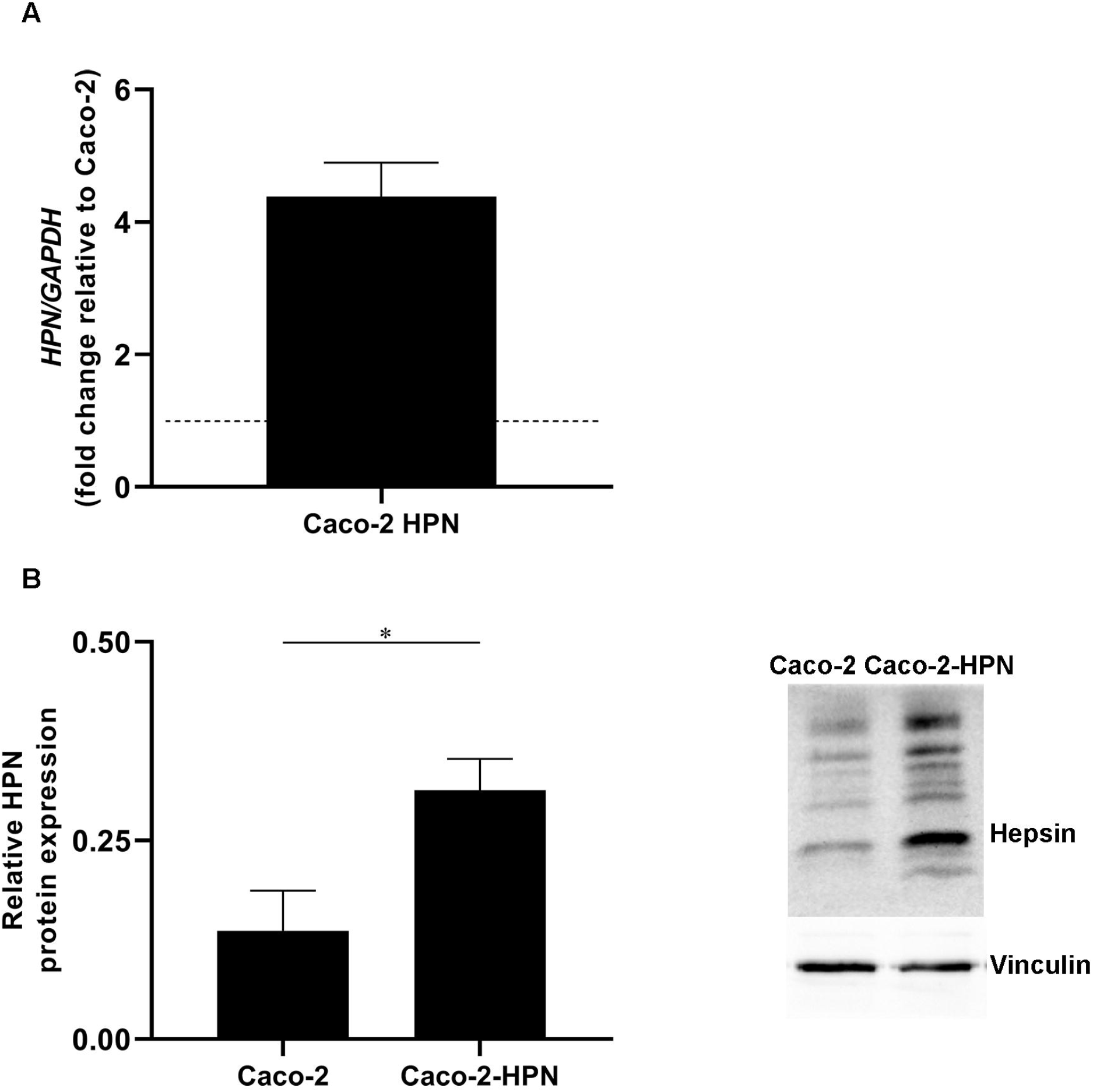
Hepsin overexpression in Caco-2-HPN cells. A) The hepsin mRNA levels were determined in Caco-2 and Caco-2-HPN cells by RT-qPCR. Gene expression levels were normalized to b-actin mRNA levels and the data are represented as the mean ± SEM of technical triplicates. Levels were shown as fold increase relative to the mean of Caco-2 cells. B) Hepsin protein levels were determined in Caco-2 and Caco-2-HPN cells by electrophoresis and western blot in lysates of Caco-2 and Caco-2-HPN cells. Vinculin expression was detected as loading control.

### 3.3. Hepsin levels affect Caco-2 cell migration and matrix degradation, but not proliferation

As enteropeptidase (another TTSP) expression regulates glioblastoma cell migration [23], we sought to probe the effect of hepsin on CRC cells. Despite the fact that these cells stand out for their reduced migration, the experiments are compatible with a slight increase in Caco-2-HPN cell migration *versus* Caco-2 cells (46·35% ± 4·87 *vs* 56·31% ± 9·99), although the difference lacks statistical significance **(Figure 3A).**

**Figure 3.**
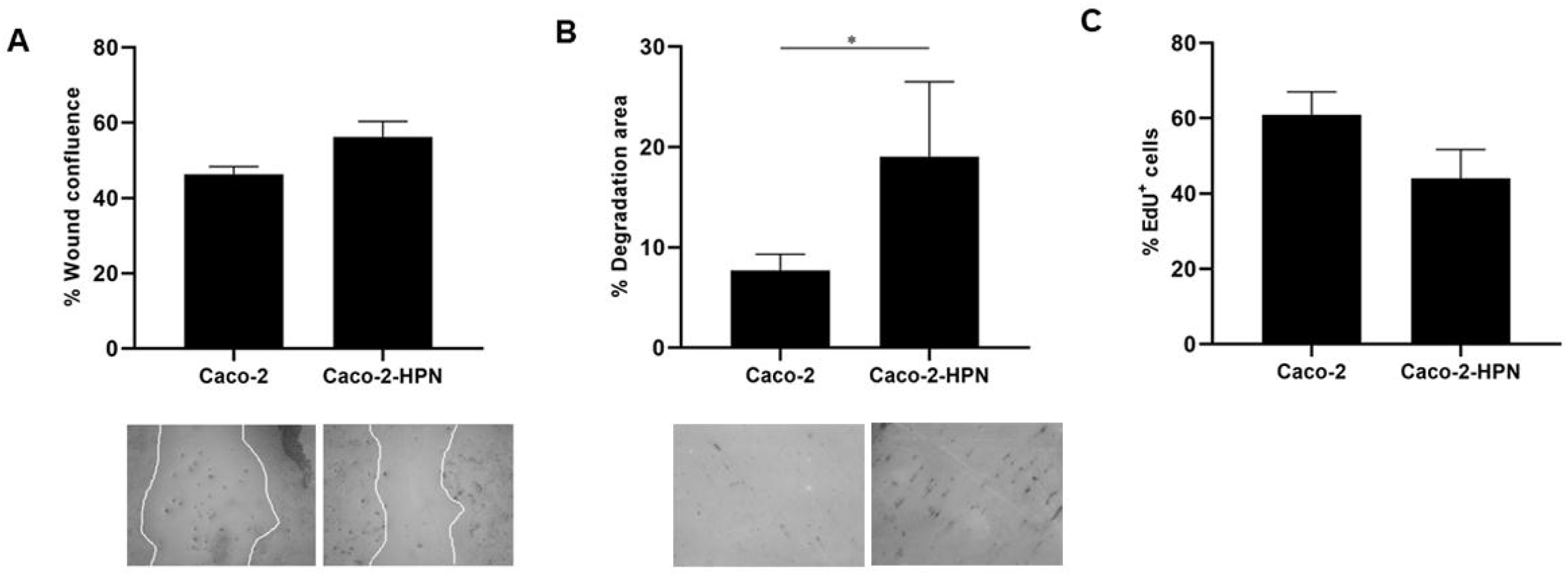
Effects of hepsin levels on cell migration, invasion, and proliferation in Caco-2 and Caco-2-HPN cells. A) Percentage of wound confluence was evaluated after 48 hours of the wound created with a pipette tip in Caco-2 and Caco-2-HPN. Six different images were processed for each sample. Images were recorded with a Leica microscope at 5× and Fiji-ImageJ was used to analyze migration. B) Percentage of cells invading were evaluated after 72 hours of matrix degradation. Images were taken with a confocal spectral scanning microscope SP8 LEICA, analyzed with ImageJ and GIMP software, and processed with Fiji-ImageJ. Cells that degraded gelatin were scored as positive. C) The percentage of total EdU+ cells were determined in Caco-2 and Caco-2-HPN cells by flow cytometry after 48h of cell culture. Each condition was evaluated in triplicate; * p<0.05; ** p<0.01; ***p<0.001.

Since hepsin is a serine protease, we also analyzed cell invasion by examining cells’ capacity to degrade a gelatin matrix. Interestingly, overexpression of hepsin significantly increased the ability of the cells to degrade gelatin compared to the basal expression (19·04 ± 8·28 vs 8·03 ± 5·05) (**Figure 3B**).

Additionally, we assessed the effect of hepsin on cell proliferation by EdU assay. In this case, there was no indication that hepsin overexpression increased proliferation (**Figure 3C**).

### 3.4. Protumor hepsin signaling pathway

Amplification of signaling mediated by extracellular signal-regulated kinases 1 and 2 (Erk1/2) has been shown to favor hepatic metastases of CRC. We therefore sought to explore the influence of hepsin expression on this pathway. **Figure 4A** shows that in Caco-2 cells, hepsin overexpression was associated with higher Erk1/2 phosphorylation (2.2 times greater than basal expression). Likewise, hepsin overexpression resulted in greater phosphorylation of signal transducer and activator of transcription 3 (STAT3) (2.2 times greater than basal expression) (**Figure 4B)**. However, despite the fact that cells with hepsin overexpression exhibited higher Akt phosphorylation, the result was not statistically significant (1.2 times greater than basal expression) (**Figure 4C**).

**Figure 4.**
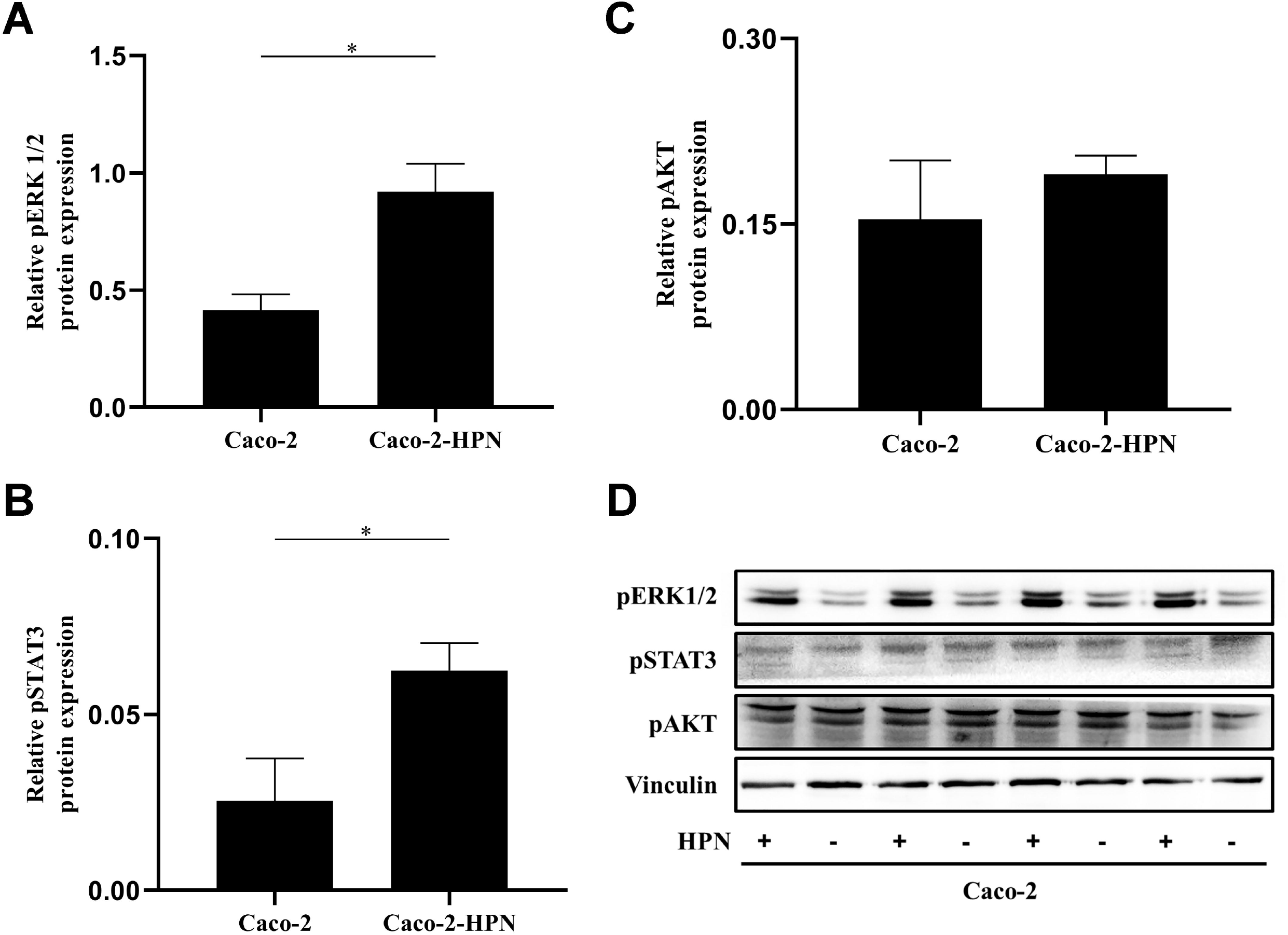
pSTAT3, pAKT, and pERK1/2 expression in Caco-2 and Caco-2-HPN cells. Expression of pERK1/2 (A), pSTAT3 (B) and pAKT (C) determined by electrophoresis and western blot in lysates of Caco-2 and Caco-2-HPN cells in triplicates. Levels were determined by densitometry and represented as relative protein expression to vinculin. D) Electrophoresis and western blot of pSTAT3, pAKT, and pERK1/2 in lysates of Caco-2 and Caco-2-HPN cells in triplicates. Vinculin expression was detected as loading control.

### 3.5. Thrombin generation assay

We then asked if high hepsin levels could contribute to generating a procoagulant state and, hence, greater thrombotic risk. We therefore performed a thrombin generation test with plasma from healthy subjects incubated with Caco-2-HPN and Caco-2 cells. As shown in both **Figure 5** and **Table 2**, exposure to Caco-2-HPN was associated with increased ETP and a higher thrombin peak and MRI compared to cells with basal hepsin expression. Moreover, we observed shorter lag time and time-to-peak in the Caco-2-HPN with respect to Caco-2 that were statistically significant.

**Figure 5.**
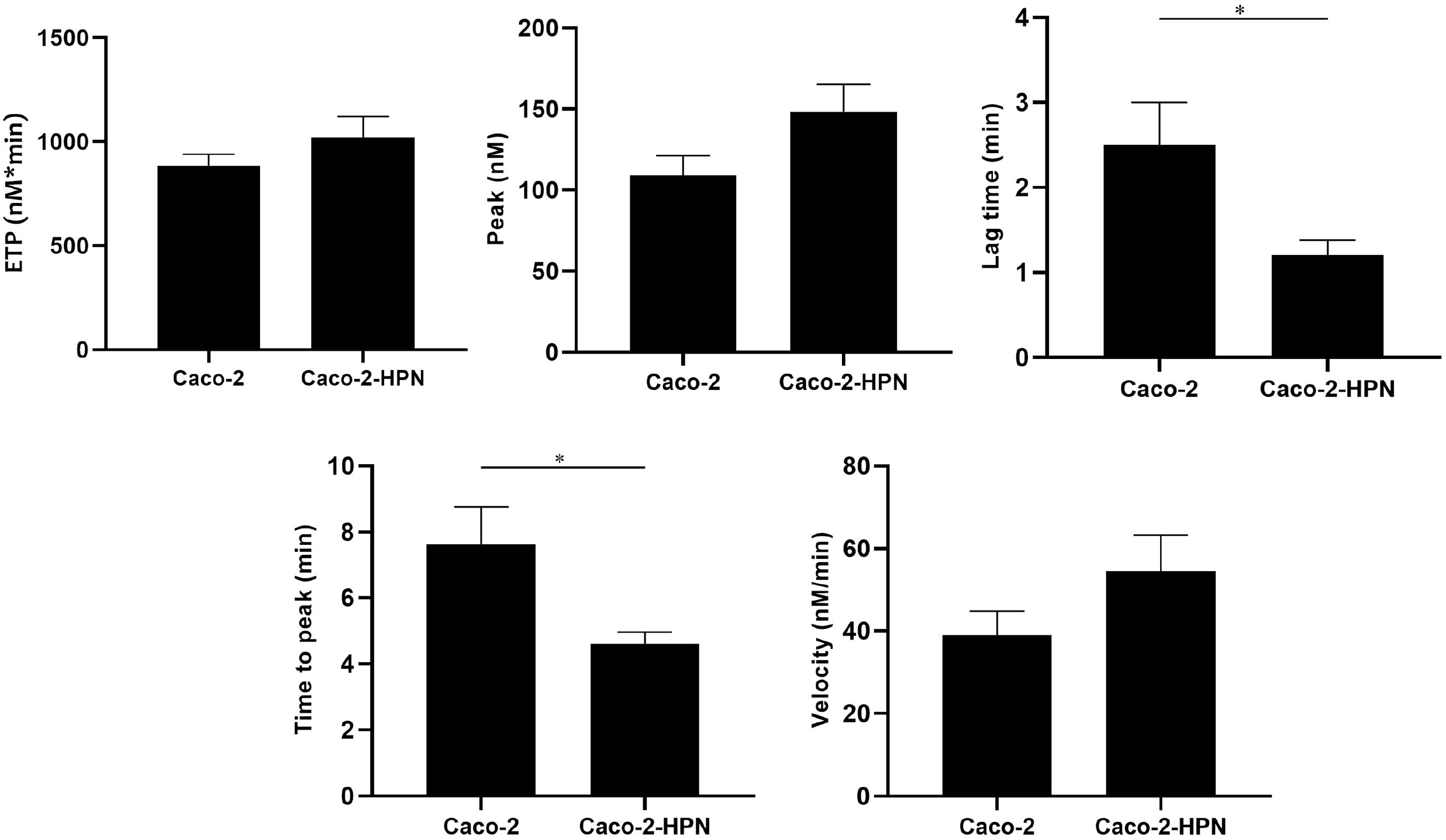
Effects of hepsin levels on thrombin generation by Caco-2 and Caco-2-HPN cells. Thrombin generation was performed after incubation of plasma with cells for 3h as described in Material and Methods. Afterwards, plasma was incubated with PPP reagent ® (final concentrations: tissue factor, 1 pmol/L; phospholipids, 4 μmol/L) and calcium chloride. The endogenous thrombin potential (ETP, nmol*min), thrombin peak (peak, nmol), lag time (min), time-to-peak (min), and mean rate index (Velocity: nmol/min) were recorded. The data represent the mean□±□SEM of at least six separate experiments. The asterisks denote statistically significant differences after Student’s t-test or the Mann-Whitney U test. *p□<□0.05.

**Table 2.**
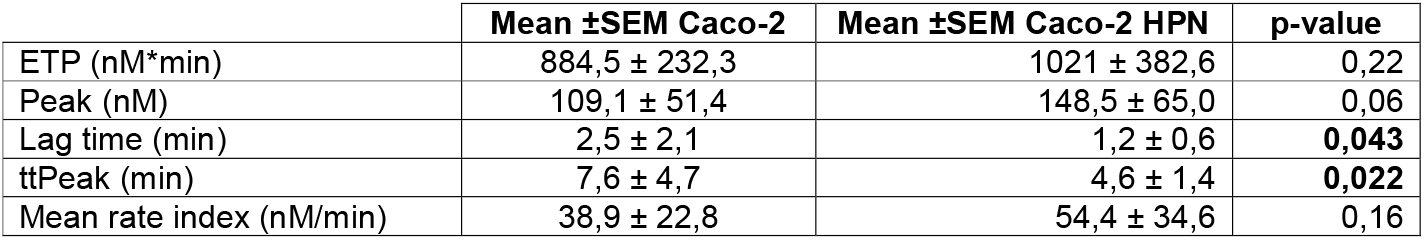
Thrombin generation assay of basal and hepsin overexpressing Caco-2 cells. P-value was calculated by unpaired t-test. Abbreviations. SEM, standard error of mean; ETP, endogenous thrombin potential; ttPeak, time-to-peak.

### 3.6. Virtual screening

Once VS calculations were completed, compounds were ranked according to their docking score, and the top 8 compounds were retained for posterior visual analysis. Venetoclax (DrugBank code DB11581) was selected and prioritized with a docking score of −11 Kcal/mol. Additional selection criteria were the hydrophobic interactions with residues PRO60, LEU41, and GLN73, and its hydrogen bond with ASN143, as illustrated in **Figure 6A**, where venetoclax impedes access to the catalytic triad, shown in surface representation for its residues.

**Figure 6.**
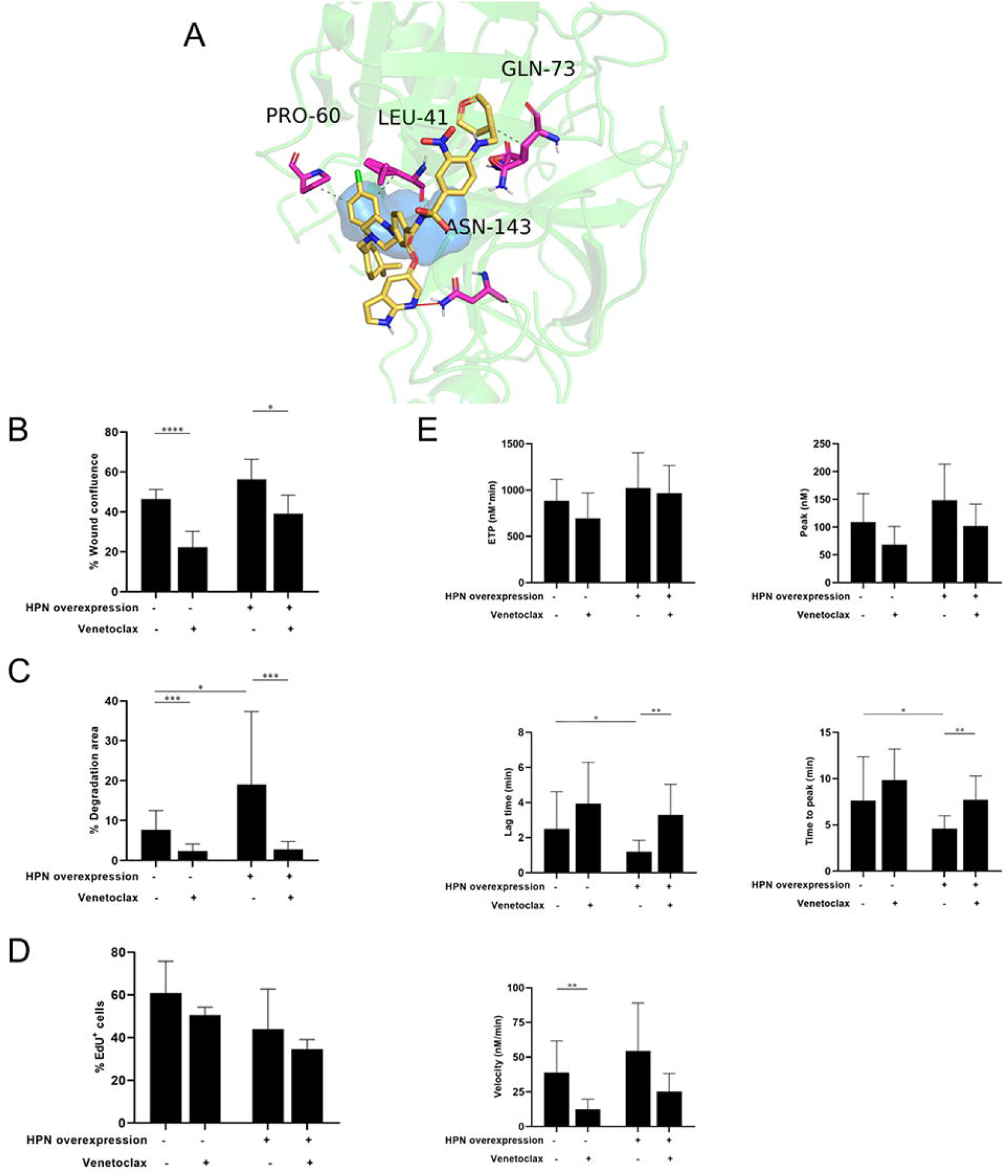
Venetoclax effect on hepsin. A) Obtained pose from molecular docking between venetoclax and hepsin. Venetoclax is shown in yellow skeleton, while hydrophobic interactions are represented in dashes, and hydrogen bond with a red line. Catalytic triad is shown in blue surface. B) Percentage of wound confluence were evaluated after 48 hours of the wound created with a pipette tip in Caco-2 and Caco-2-HPN, in the presence and absence of 1.88 μM venetoclax. Six different images were processed for each sample. Images were recorded with a Leica microscope at 5× and Fiji-ImageJ was used to analyze invasion. C) Percentage of cells invading were evaluated after 72 hours of matrix degradation in presence and absence of 1.88 μM venetoclax. Images were taken with a confocal spectral scanning microscope SP8 LEICA, analyzed with ImageJ and GIMP software, and processed with Fiji-ImageJ. Cells that degraded gelatin were scored as positive. D) The percentage of total EdU+ cells were determined in Caco-2 and Caco-2-HPN cells by flow cytometry after 48h of cell culture in presence and absence of 1.88 μM venetoclax. E) Thrombin generation was performed after incubation of plasma with cells in presence and absence of 1.88 μM venetoclax as described in Material and Methods. The endogenous thrombin potential (ETP, nmol*min), thrombin peak (peak, nmol), lag time (min), time-to-peak (min), and mean rate index (velocity: nmol/min) were recorded. The data represent the mean ± SEM of at least six separate experiments. The asterisks denote statistically significant differences after Student’s t-test or the Mann-Whitney U test. Each condition was evaluated in triplicate; * p<0.05; ** p<0.01; ***p<0.001.

### 3.7. Venetoclax inhibition of hepsin

We tested the effect of venetoclax on hepsin by evaluating the proteolytic activity of hepsin on a fluorogenic substrate. Venetoclax was capable of irreversibly inhibiting hepsin activity. We therefore calculated the half maximal inhibitory concentration (IC50) by incubating different concentrations of venetoclax and monitoring hepsin activity. As shown in supplementary **Figure S3**, the IC50 of venetoclax was 0.48 μM. This value was higher for hepsin than for the one recently reported for indole derivatives [33]. The toxicity of these compounds has yet to be tested in humans, while the calculated IC50 of venetoclax for hepsin is within the concentration range tested in chronic lymphocytic leukemia cells [34].

### 3.8. Venetoclax effect on CRC cells

Venetoclax significantly reduced cell migration both in cells with basal expression (46·35% ± 4·87 *vs* 22·26% ± 7·97), as well as in cells that overexpress hepsin (56·31% ± 9·99 *vs* 39·09% ± 9·27) (**Figure 6B**). As **Figure 6C** illustrates, venetoclax was able to specifically inhibit Caco-2 and Caco-2-HPN cell invasion to a statistically significant degree (2·08% ± 1·81 vs 8·03% ± 5·05 and 2·98% ± 2·03 vs 19·04% ± 18·28, respectively). Despite venetoclax being a Bcl-2 inhibitor, we did not find evidences of venetoclax treatment affecting CRC cell proliferation or cell cycle (**Figure 6D, Figure S4**). Finally, incubating the cells with venetoclax led to a statistically significant reduction in the rate of thrombin generation velocity in the cells with basal expression and prolonged lag time and time-to-peak in both cell lines (**Figure 6E, Table 3**). This effect was not shown in cells without hepsin expression, such as DLD1 and HCT-116 (**Figure S5**, **Figure S6** and **Supplementary table 1**).

**Table 3.** Thrombin generation assay of basal and hepsin overexpressing Caco-2 cells in the presence and absence of venetoclax. P-value was calculated by unpaired t-test. Abbreviations. SEM, standard error of mean; ETP, endogenous thrombin potential; ttPeak, time-to-peak.

| | Mean ±SEM | | | | p-value | |
| --- | --- | --- | --- | --- | --- | --- |
|  | Caco-2 | Caco-2 + Venetoclax | Caco-2 HPN | Caco-2 HPN + Venetoclax | Caco-2 vs Caco-2 + Venetoclax | Caco-2 HPN vs Caco-2 HPN + Venetoclax |
| ETP (nM*min) | 884,5 ±232,3 | 697,3 ±271,9 | 1021 ±382,6 | 967,1 ±299 | 0,14 | 0,78 |
| Peak (nM) | 109,1 ±51,4 | 68,71 ±32,5 | 148,5 ±65 | 101,7 ±39,7 | 0,11 | 0,15 |
| Lag time (min) | 2,5 ±2,1 | 3,9 ±2,4 | 1,2 ±0,6 | 3,3 ±1,7 | 0,21 | **0,001** |
| ttPeak (min) | 7,6 ±4,7 | 9,8 ±3,4 | 4,6 ±1,4 | 7,7 ±2,6 | 0,35 | **0,003** |
| Mean rate index (nM/min) | 38,9 ±22,8 | 12,4 ±7,4 | 54,4 ±34,6 | 25,2 ±13 | **0,022** | 0,09 |

### 3.9. Venetoclax effect on CRC cells invasiveness in vivo

The higher *in vitro* invasiveness of Caco-2-HPN was also confirmed *in vivo* using a xenotransplantation model in zebrafish larvae. More interestingly, pre-treatment of these cells with venetoclax reduced its invasiveness to the levels found in parental cells (**Figure 7**), confirming the *in vitro* studies and further showing its therapeutical potential for the treatment of CRC.

**Figure 7.**
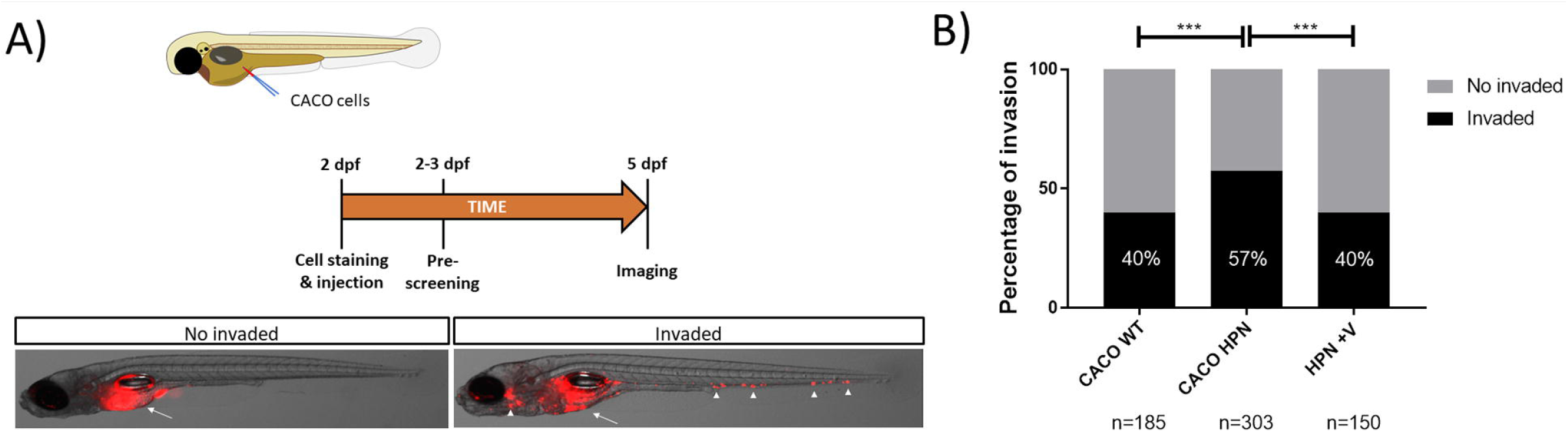
Venetoclax reduces Caco-2-HPN cell invasion in zebrafish larvae. A) Experimental design of the zebrafish larvae xenotransplantation experiments and representative pictures of no invaded and invaded larvae showing red-labelled Caco-2 cells at the injection site (arrow) and several invasion foci (arrowheads). B) Evaluation of invasion score 3 days after injection in the yolk sac of 2 dpf casper zebrafish larvae. The results showed are a pool of three different experiments. ***p < 0.001 according to a χ^2^ test. n, number of larvae per treatment. V, venetoclax.

## 4. Discussion

Hepsin is one of the TTSPs that exert a complex influence on different kinds of oncogenic processes [3–6,10–12], parallel to the activation of the hemostatic system, and that as such, comprise critical nodes where multiple signaling pathways converge with coagulation cascade effectors. CRC is a suitable model to explore these interactions due to the downstream regulation of hepsin dependent on the Ras/Erk1/2 signaling pathway [7], as well as the worse prognosis and greater thrombotic risk associated with the activation of this signaling pathway [35,36]. Information about the role of hepsin in CRC is exceedingly scant. Recently, we have identified hepsin as a prognostic marker of metastasis and thrombosis in patients with localized tumor [13]. This led us to explore possible hepsin-mediated mechanisms that might link to CRC tumorigenesis and thrombotic risk, as well as to search for drugs that target this TTSP. In addition, although hepsin’s serine protease domain is located in the extracellular region, there are reports that support it can also be secreted, making it possible to monitor it in blood [37]. This enables us to evaluate it as a possible prognostic or predictive biomarker of thrombotic risk in plasma.

First of all, what we have found here is that plasma hepsin levels are higher in individuals with active metastatic disease compared to patients who have undergone tumor resection. This corroborates the findings of Tieng et al., who have reported the presence of elevated serum hepsin values in subjects with metastatic CRC [9], which prompts it to be proposed as a serum/plasma biomarker. Though the statistical evidence is lower, our results are also consistent with earlier observations that implicate hepsin in processes related to Ras-dependent tumorigenesis [7]. However, these results are different from those found by our group in the biopsies of patients with CRC [13]. In the primary tumor, hepsin appears to play an initiating role in metastasis. However, once metastasis has started, the role of hepsin in the primary tumor is less clear. Hepsin has an extracellular fraction that can be released from the transmembrane region and remain in plasma. The fact that hepsin is differentially more expressed in the plasma of metastatic patients could be due to its release by the cells from the metastasis sites and not from the primary tumor, where the role of hepsin would no longer be as relevant. In fact, hepsin levels in the primary tumor are lower the greater the number of metastases is [13].

Secondly, in an attempt to shed additional light on the underlying mechanism of these findings regarding tumorigenesis, we have studied the influence of hepsin on a CRC cell line. Our results confirm that hepsin expression is associated with greater Caco-2-HPN cell migration and invasion. This is in line with findings in other pathologies such as prostate cancer, in which hepsin regulates cell migration by proteolyzing laminin-332, a cell matrix molecule necessary for cell-to-cell adhesion, thereby enhancing tumor cell invasion and migration [38]. These actions are exerted directly or by means of inducing and activating matrix metalloproteinases, resulting in a potent destruction of the extracellular matrix [39]. Indeed, hepsin is not the only TTSP involved in CRC invasion and metastasis. The up-regulation of other key TTSPs, such as tissuetype plasminogen activator, urokinase-type plasminogen activator, plasminogen activator inhibitor type-1, and others, play an essential role in CRC tumorigenesis [40]. Together, they all comprise a complex, redundant, multi-stage system that, as a whole determines the degradation of the basement membrane and extracellular matrix by proteinases. In turn, these actions constitute a stimulus for the hemostatic system.

Moreover, we have found that hepsin overexpression is associated with greater Erk1/2 phosphorylation and greater STAT3 activation. Erk1/2-mediated signaling amplification has been proven to promote hepatic metastases in CRC [41], which accounts for the higher hepsin levels in advanced tumors. Interestingly, Erk1/2 activation, but not Akt activation, predicts worse prognosis in CRC [42]. At the same time, the JAK2/STAT3 signaling pathway plays an important role in regulating apoptosis and enhancing clonogenic potential [43]. Nevertheless, we do not know the mechanism by which hepsin regulates the activation of these pathways. This is a matter that merits study in the future. Hepsin is capable of cleaving the epidermal growth factor receptor (EGFR), so that the fragments are tyrosine-phosphorylated, but it is not clear which downstream signaling pathways are activated as a result. In fact, hepsin-induced EGFR cleavage does not go hand-in-hand with increased Erk1/2 [44,45]. Instead, hepsin activity is regulated downstream via a Ras pathway-dependent mechanism [7]. However, in our recently published study, we found no association between Ras mutation and hepsin levels [13].

The proteolytic enzymes involved might also account for the two-way links between thrombosis and cancer [20,46], in the degree to which the degradation of the extracellular matrix by proteinases is coupled to the activation of the hemostatic system. The incidence of thromboembolic disease was low in our plasma series, in keeping with the bibliographic data [47,48]. Despite the evidence being anecdotal, it is nonetheless striking that none of the individuals with hepsin levels <Q3 exhibited a thrombotic event in the first 24 months. Interestingly, in our recently published study, hepsin is a potential biomarker of thrombosis in patients with localized CRC [13]. The fact that hepsin is not associated with increased thrombotic risk in biopsies from metastatic patients could be due to its detection in plasma, where it would have more access to coagulation proteins. In fact, thromboembolic disease is often associated with metastasis [46]. Hepsin overexpression of our cell line clearly increased procoagulant phenotype as we observed in localized CRC patients [13]. These observations led us to hypothesize that some TTSPs, including hepsin, contribute to hypercoagulability *in vivo* by generating thrombin in the phase boundary of the tumor invasion and migration front, together with other mediators [49,50]. In the future, it would be interesting to study whether there is an inverse correlation between hepsin levels in the biopsy and in the plasma of metastatic patients to corroborate these hypotheses.

Third, we also wanted to probe into hepsin’s role as a potential therapeutic target for CRC. Initially we carried out the silencing of hepsin expression using siRNAs (**Figure S1**), however, because the migration and invasion experiments took 72 hours the transient silencing was reversed and we did not observe differences of the silenced cells with Caco-2 and Caco-2-HPN cells. Therefore, we decided to carry out a different strategy. To do so, we used VS to identify potential hepsin inhibitors. We identified venetoclax, a drug used in combination to treat acute myeloid leukemia and relapsed or refractory chronic lymphocytic leukemia [51,52]. We have demonstrated that venetoclax is a specific inhibitor of all hepsin’s protumoral and prothrombotic actions identified in this study in CRC cells. It is important to note that venetoclax also exerts these inhibitory effects on Caco-2 cells because these cells also have a basal hepsin expression. In fact, venetoclax exerts no so clear effects on thrombin generation in colorectal cancer cells without hepsin expression (**Figure S5 and supplementary Table 1**). It is surprising that despite venetoclax being a BCL2 inhibitor, an antiapoptotic protein that is pathologically overexpressed and crucial to the survival of certain cancer cells, Caco-2 cell proliferation nor its cell cycle is affected by treatment with this drug. This may be due to the existence of drug efflux pumps that prevent it from reaching the inside of the cell [53], although this has not been investigated in this study. This might also account for the lack of effect venetoclax has on intracellular signaling pathways that could be independent of the activity of serine proteases. Overall, these observations support the notion that the antitumor effect of venetoclax in Caco-2 cells takes place in the extracellular space by inhibiting hepsin. Most importantly, venetoclax significantly reduces invasion of hepsin-overexpressing cells to levels found in parental cells, showing its efficacy in an *in vivo* model. While other hepsin inhibitors have been reported, they are limited by their decreased stability *in vivo*. Moreover, some have off-target actions, diminishing their effectiveness on hepsin [54].

Undoubtedly, our study has limitations the reader should bear in mind. First of all, the scant events do not allow for a solid analysis of the association between plasma hepsin and thrombotic risk, as well as survival endpoints. As with any other biomarker, interpreting the effect of its levels is complex, as they are contingent on specific molecular pathways (e.g., the mitogen-activated protein kinase pathway) and presumably depend on circumstances such as tumor burden, tumor response to chemotherapy, or antagonistic or paradoxical effects [11,14,15]. Consequently, the results should be interpreted as hypothesis generators, to be reviewed later. In addition, there is no correlation between the findings found in plasma and biopsies of patients with CRC [13], so more studies are needed to determine the potential of hepsin as a biomarker and as a therapeutic target. Second, although the study sheds light on the status of two key pathways in CRC (Erk1/2 and JAK2/STAT3 signaling pathways), the causal order is unclear in this case. Our study provides an overall panorama to be supplemented later on [7]. Third, our research reveals the antitumor and potentially antithrombotic effect of venetoclax, an FDA-approved oral agent, in an *in vivo* model of CRC possibly due to anti-hepsin activity, although it cannot be ruled out that this effect might be mediated by its action on other proteins. It would be also necessary to identify those patients with CRC who could benefit from the effect of venetoclax in a clinical trial. Finally, the reader should be mindful of the tremendous clinical heterogeneity of CRC that may have affected these results.

## 5. Conclusions

In short, our results demonstrate that high hepsin levels may be indicative of a more aggressive tumor phenotype, because of the higher potential for migration and invasion in colorectal cancer cells and greater activation of coagulation. Furthermore, our study provides hepsin as a novel therapeutic target against these malignancies and their underlying pathologies. Finally, this study reports, for the first time, an anti-migratory, anti-invasive, and potentially anti-thrombotic effect of venetoclax, which could indicate its use as a new molecular targeted treatment in CRC and, potentially, other hepsin-overexpressing tumors.

## Data accessibility

The original data are available upon request to the authors.

## Author contributions

Conceptualization, formal analysis, funding acquisition, project administration, supervision, writing: I.M.M., H.P.S. and A.C.B.

Investigation, methodology and data curation: M.C.R., J.P., I.P.S, D.Z.H., C.O., S.E., G.L.G, J.P.G., G.R., S.M., F.A., F.B., F.G.M., and A.N

Original draft and writing: M.C.R.

Review & editing: F.B., V.V., and V.M

HPS, ACB, and IMM had full access to data; HPS, ACB and IMM were ultimately responsible for the decision to submit this manuscript for publication.

## Acknowledgements

The authors wish to thank María Eugenia de la Morena-Barrio, Javier Corral, Manuel Sánchez-Canovas, and Alberto Martínez for their technical assistance.

This work was funded by the Carlos III Health Institute (ISCIII) (Grant Number PI17/00050 & FEDER, PI21/00210 & FEDER), Spanish Ministry of Economy and Competitiveness (Grant numbers: BIO2017-84702-R & FEDER and CTQ2017-87974-R), Fundación Séneca (Project 20988/PI/18) and the generous donations to Crowdfunding Precipita (FECYT).

Supercomputing resources in this work have been supported by the Poznan Supercomputing Center’s infrastructures, the e-infrastructure program of the Research Council of Norway, and the supercomputing center of UiT—the Arctic University of Norway, by the Plataforma Andaluza de Bioinformática of the University of Málaga, by the supercomputing infrastructure of the NLHPC (ECM-02, Powered@NLHPC), and by the Extremadura Research Centre for Advanced Technologies (CETA-CIEMAT), funded by the European Regional Development Fund (ERDF). CETA-CIEMAT belongs to CIEMAT and the Government of Spain.

Maria Carmen Rodenas has a grant from the Spanish Society of Thrombosis and Haemostasis (SETH), David Zaragoza-Huesca holds a predoctoral FPU grant from the Spanish Ministry of Science and Innovation, Ginés Luengo-Gil has a grant from the Spanish Society of Hematology and Hemotherapy and Irene Martínez-Martínez holds a Miguel Servet contract (CPII18/00019) from the Carlos III Health Institute (Madrid).

The funders had no role in study design, data collection, analysis and interpretation, or manuscript writing and submission.

## Conflict of Interest

This work is included in a patent application.

## Abbreviations

CI: cumulative incidence
CRC: colorectal cancer
DAPI: 4’, 6-diamino-2-phenylindole
EdU: 5-ethynyl-2’-deoxyuridine
EGFR: epidermal growth factor receptor
Erk1/2: extracellular signal-regulated kinases
ETP: endogenous thrombin potential
*HPN*: hepsin gene
IC50: half maximal inhibitory concentration
KRAS: Kirsten Rat Sarcoma Viral Oncogene Homologue
STAT3: signal transducer and activator of transcription 3
TTSPs: type II transmembrane serine proteases
VS: virtual screening

## Notes

### Competing Interest Statement

The authors have declared no competing interest.

