## Supplemental information for "Pro-tumor and prothrombotic activities of hepsin in colorectal cancer cells and suppression by venetoclax"

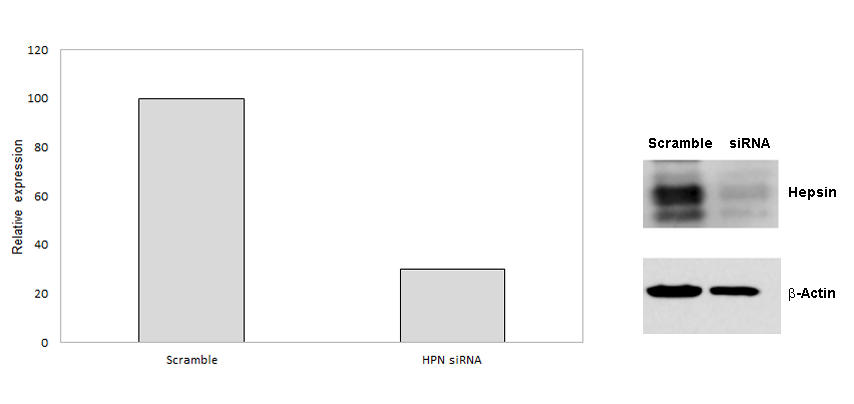

**Figure S1. *HPN* gene silencing efficiency in Caco-2 cells.** Hepsin protein levels were determined by electrophoresis and western blot in lysates of Caco-2 transfected with scramble or 5 nM ON-TARGETplus SMARTpool siRNAs against *HPN*. Beta-actin expression was used as a loading control.

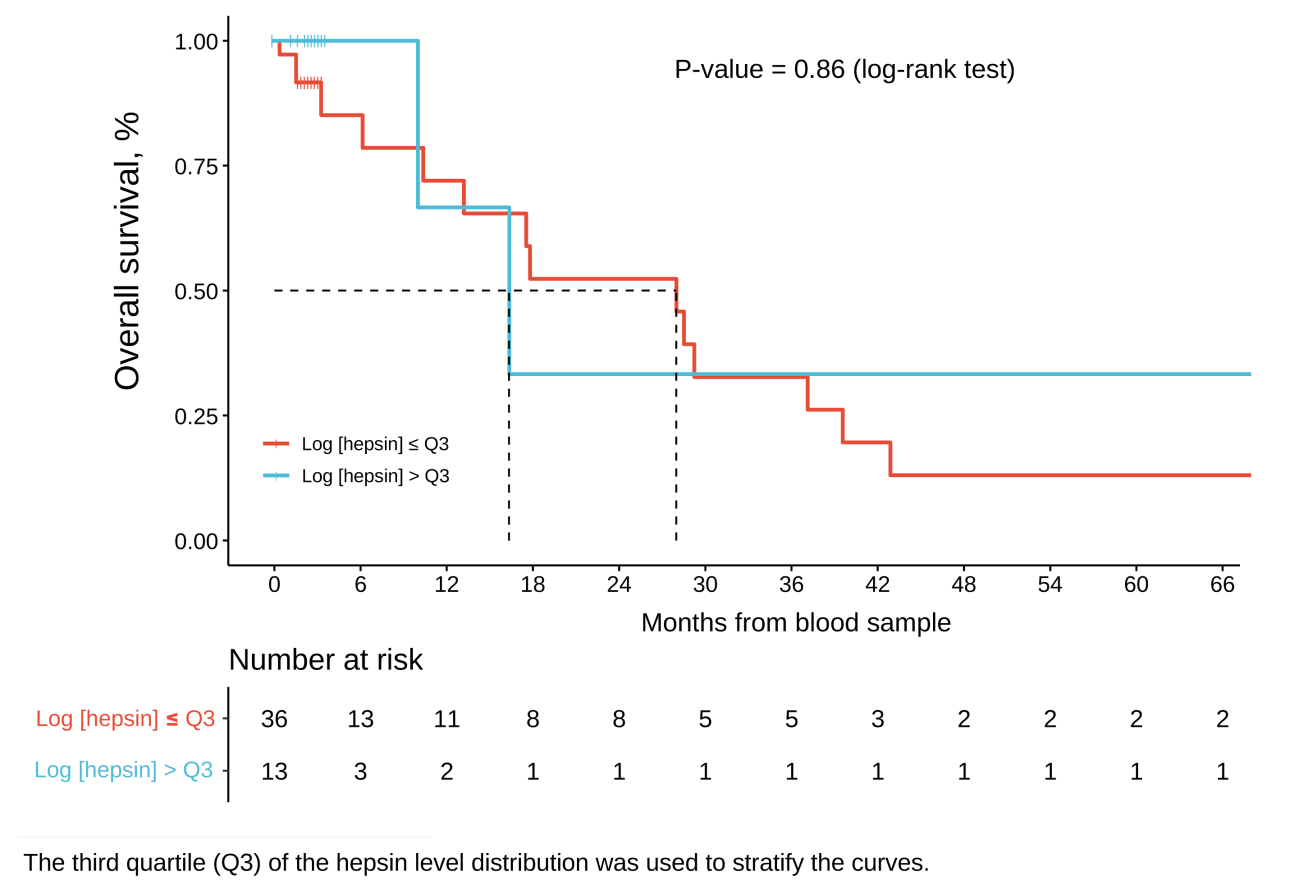

**Figure S2. Kaplan-Meier plots for overall survival based on hepsin blood levels.**

**Figure S3. Calculated IC50 value of Venetoclax on the activity of hepsin.** Maximum activity rate of hepsin in the presence of increasing concentrations of Venetoclax. FU: Fluorescence Units.

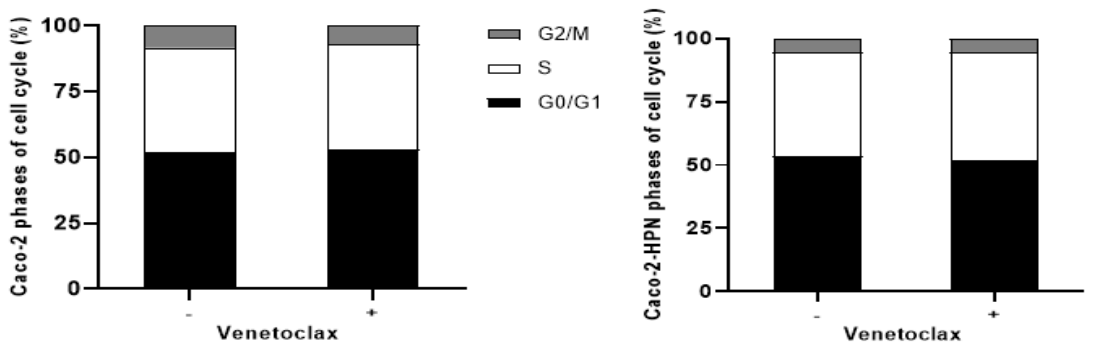

**Figure S4. Effect of hepsin expression and venetoclax on cell cycle distribution.** Cell cycle distribution was analyzed by flow cytometry and representes as a histogram. Data are representative of at least three independent experiments.

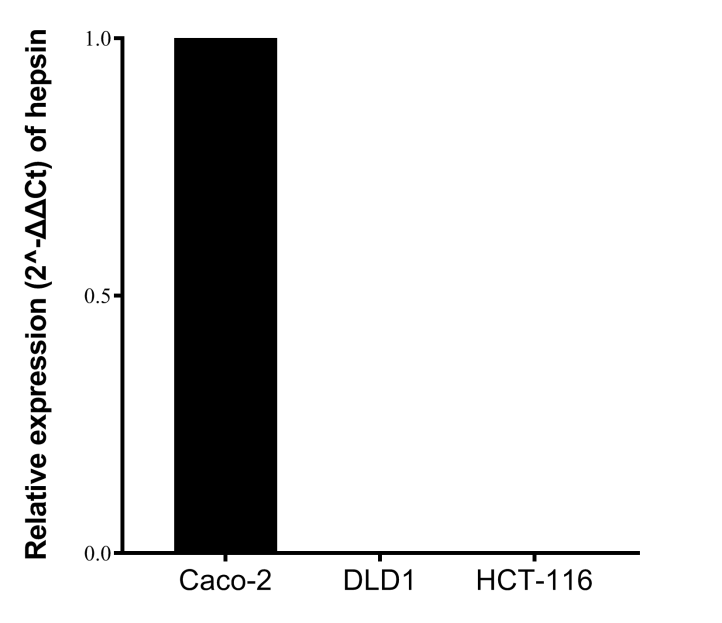

**Figure S5.** **Relative expression (2-ΔΔCT) of DLD1 and HCT-116 cell lines compared to relative Caco-2 expression.**

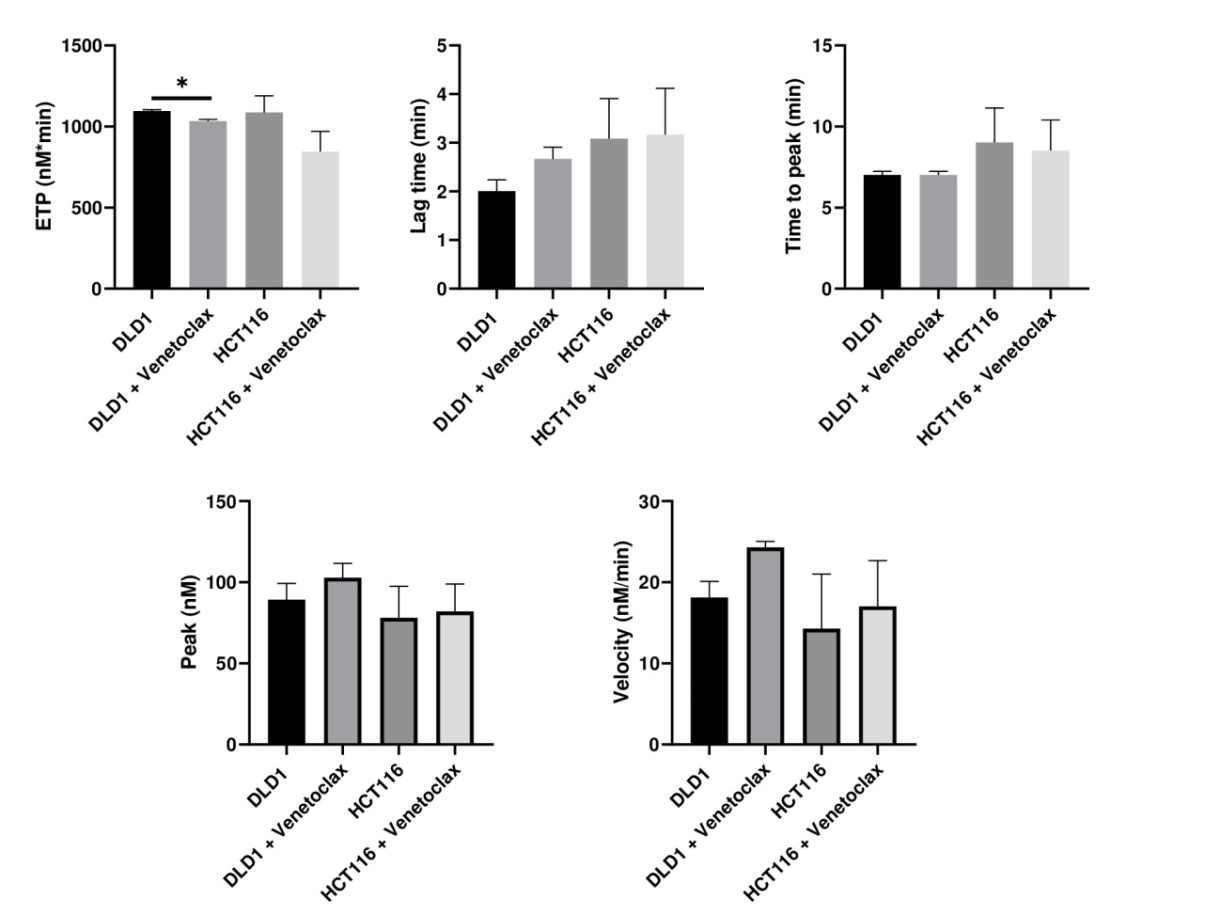

**Figure S6.** **Effects of Venetoclax on thrombin generation by DLD1 and HCT116 cells.** Thrombin generation was performed after incubation of plasma with cells for 3h as described in Material and methods. Afterwards, plasma was incubated with PPP reagent ® (final concentrations: tissue factor, 1 pmol/L; phospholipids, 4 μmol/L) and calcium chloride. The endogenous thrombin potential (ETP, nM*min), thrombin peak (peak, nM), lag time (min), time to peak (min), and mean rate index (Velocity: nM/min) were recorded. The data represent the mean ± SEM of at least four separate experiments. The asterisks denote statistically significant differences after Student’s t-test. ***: p < 0.05.

**Supplementary table 1.** Thrombin generation assay of DLD1 and HCT116 cells in the presence and absence of venetoclax. P-value was calculated by Student´s t-test. *SEM*: standard error of mean; *ETP*: endogenous thrombin potential.

|  | **Mean ±SEM** | | | | **p-value** | |
| --- | --- | --- | --- | --- | --- | --- |
|  | DLD1 | DLD1 + Venetoclax | HCT116 | HCT116 + Venetoclax | DLD1 vs DLD1 + Venetoclax | HCT116 vs HCT116 + Venetoclax |
| **ETP (nM*min)** | 1095.695±8.309 | 1032.665±12.325 | 1087.405±101.901 | 845.39±124.917 | **0.0267** | 0.1677 |
| **Peak (nM)** | 89.330±10.041 | 102.835±8.846 | 78.020±19.516 | 82.075±16.907 | 0.2897 | 0.8449 |
| **Lag time (min)** | 2.005±0.233 | 2.67±0.240 | 3.085±0.827 | 3.17±0.948 | 0.1069 | 0.9326 |
| **Time to peak (min)** | 7.015±0.233 | 7.015±0.233 | 9.015±2.128 | 8.515±1.888 | >0.9999 | 0.8269 |
| **Mean rate index (nM/min)** | 18.115±2.001 | 24.310±0.721 | 14.295±6.725 | 17.03±5.643 | 0.0542 | 0.7025 |
